# Mitochondrial alternative oxidase contributes to successful tardigrade anhydrobiosis

**DOI:** 10.1101/2020.09.25.313122

**Authors:** Daria Wojciechowska, Andonis Karachitos, Milena Roszkowska, Wiktor Rzeźniczak, Robert Sobkowiak, Łukasz Kaczmarek, Jakub Kosicki, Hanna Kmita

## Abstract

Anhydrobiosis can be described as an adaptation to lack of water. This adaptation provides some organisms including tardigrades with a set of capabilities allowing them to survive extreme conditions that even do not exist on Earth. However, the underlying cellular mechanisms are still not explained. Available data assumes important contribution of mitochondrial proteins. Since mitochondrial alternative oxidase (AOX) described as a drought response element has recently been proposed for various invertebrates including tardigrades, we have decided to check if AOX is involved in successful anhydrobiosis of tardigrades. *Milnesium inceptum* was used as a model for the study. We confirmed functionality of *M. inceptum* AOX and estimated its activity contribution to anhydrobiosis of different duration. We observed that AOX activity was particularly important for *M. inceptum* revival after longer-term anhydrobiosis but did not affect rehydration stage. The results may contribute to explanation and then application of anhydrobiosis underlying mechanisms.

## INTRODUCTION

It is truism to write that water availability is one of the most important factors for life. However, terrestrial habitats may endure occasional lack of water that requires from inhabiting organisms specific adaptations to drought, functioning at different levels of biological organization. One of the most prevalent adaptations is anhydrobiosis, often called simply “life without water” (e.g. Watanabe, 2006; Rebecchi, 2013; Arakawa and Blaxter, 2017; Kaczmarek et al., 2019). The phenomenon is also known as desiccation tolerance that in turn is defined as the ability to dry to equilibrium with moderately to very dry air (i.e. to 10% water content or even less) and then to recover to normal functioning after rehydration without sustaining damages (Alpert, 2006). Explanation of mechanisms underlying anhydrobiosis and consequently indication of successful anhydrobiosis biomarkers may contribute to different applications because of anhydrobiotic organism tolerance to extreme environmental conditions. This concerns, for example, dry vaccines, preservation of biological materials for transplantation or food production, enzymes working in a small amount of water and mechanisms of DNA protection and repair (e.g. Schill et al., 2009; Wełnicz et al., 2011; Guidetti et al., 2012; Ono et al., 2016; Erdman and Kaczmarek, 2017; Kaczmarek et al., 2019).

Anhydrobiosis has been reported for many microorganisms as well as for plants and some small invertebrates and among the latter the best known examples are tardigrades (e.g. Wright et al., 1992; Bertolani et al., 2004; Alpert, 2006; Rebecchi et al., 2007; Schokraie et al., 2010; Nelson et al., 2015; Schill and Hengherr, 2018; Kaczmarek et al., 2019). They are water-dwelling eight-legged segmented invertebrates found almost everywhere, from mountaintops to the deep sea (Nelson et al., 2015). Depending on the habitat tardigrades are defined as terrestrial, freshwater and marine ones (Nelson et al., 2018). The terrestrial tardigrades need a film of water to be active, and therefore they are also termed limno-terrestrial (also semi-terrestrial or semi-aquatic). This group of tardigrades includes most of the species undergoing successful anhydrobiosis (e.g. Wright, 1989; Hygum et al., 2016; Møbjerg et al., 2018; Sørensen-Hygum et al., 2018). Tardigrade anhydrobiosis includes entering, permanent and leaving stages corresponding to the dehydration, tun and rehydration stages, respectively (e.g. Rebecchi et al., 2007). Specifically, the tun stage is a dehydrated tardigrade that “shrunk” to about 30% of its original volume to a state resembling a barrel. This includes contraction of anterior-posterior body axis, retraction of legs, and rearrangement of internal organs and cells, and these changes are reversed during the rehydration stage (e.g. Wright et al., 1992; Rebecchi et al., 2007, 2013; Nelson et al., 2015; Møbjerg et al., 2018). Thus, on the organismal level anhydrobiosis seems to be a fairly well understood process.

However, tun formation is a coordinated process (Crowe, 1975) that requires adequate molecular mechanisms. Although searching on the mechanisms has revealed their differentiation between tardigrade species (e.g. Hengherr et al., 2008; Rebecchi et al., 2013; Bemm et al., 2017; Gross et al., 2019; Kamilari et al., 2019), available data appears to indicate common and important role of proper mitochondria functioning. Accordingly, it has been shown that mitochondria uncoupling abolishes tun formation (Halberg et al., 2013) and reactive oxygen species (ROS) generated in majority by mitochondria (e.g. Murphy, 2009) are described as signals important for adaptation to drought and desiccation (Scheibe and Beck, 2011; Rogov and Zvyagilskaya, 2015). Additionally, it is suggested that mitochondria contribute to tun functionality and successful rehydration (Pigoń and Węglarska, 1955; Jönsson and Rebecchi, 2002; Tanaka et al., 2015) but their role in tardigrade anhydrobiosis still remains elusive. Therefore, studies on mitochondrial proteins described as markers of stress conditions is a reasonable step in understanding of the role of mitochondria in anhydrobiosis.

One of the mitochondrial proteins described as the marker of stress conditions and an important element of the response to drought stress is alternative oxidase (e.g. Pastore et al., 2007; Hanqing et al., 2010; Rogov and Zvyagilskaya, 2015; Vanlerberghe et al., 2016; McDonald and Gospodaryov, 2019; Weaver, 2019). Mitochondrial alternative oxidase (AOX) is the mitochondrial inner membrane, cyanide insensitive, iron-binding protein which introduces a branch into the mitochondrial respiratory chain at the coenzyme Q level. This allows for electron transfer from the respiratory chain complexes I and II via coenzyme Q to oxygen without involvement of the respiratory chain cytochrome pathway composed of complexes III and IV (for review, see e.g. Sluse and Jarmuszkiewicz, 1998; Moore and Albury, 2008; Rogov and Zvyagilskaya, 2015; Weaver, 2019). As a result AOX provides a bypath that releases constraints on the respiratory chain cytochrome pathway and consequently participates in mitochondrial reduction-oxidation reactions important for cell metabolic plasticity involved in adaptation to variable biotic and abiotic stress factors. Accordingly, heterologous expression of AOX in cultured mammalian cells, fruit flies and mice under mitochondrial respiratory stress conditions results in restoration of respiratory activity and metabolism correction (e.g. Szibor et al., 2020).

AOX was initially considered to be limited to plant as well as some fungus and protist species but its presence has recently been proposed for different invertebrate animals, except for insects (e.g. McDonald et al., 2009; Pennisi et al., 2016; McDonald and Gospodaryov, 2019; Weaver, 2019; Szibor et al., 2020). Analysis of available but scarce tardigrade genomic and/or transcriptomic data indicates the presence of AOX encoding gene in three tardigrade species (Bemm et al., 2017; McDonald and Gospodaryov, 2019) namely, *Hypsibius exemplaris* Gąsiorek, Morek and Michalczyk, 2018 (former *Hypsibius dujardini* (Doyère, 1840)), *Ramazzottius varieornatus* Bertolani and Kinchin, 1993 and *Milensium inceptum* Morek, Suzuki, Schill, Georgiev, Yankova, Marley and Michalczyk, 2019. The latter, formerly assigned as *Milnesium tardigradum* Doyère, 1840 (Morek et al., 2019), is known to be highly resistant to periodical dehydration events under laboratory conditions (e.g. Hengherr et al., 2008). Thus, in this study we tested hypothesis that AOX is involved in tardigrade anhydrobiosis using *M. inceptum* as a model for the research. To test this hypothesis we verified functionality of *M. inceptum* AOX and then estimated the animal recovery to full activity after tun formation or rehydration in the presence of AOX inhibitor, using different duration of anhydrobiosis. The obtained results indicate that AOX activity is important for tun revival reflected by their ability to return to full activity but does not appear to affect importantly the rehydration stage itself. Additionally, the contribution of AOX depends on anhydrobiosis duration.

## MATERIALS AND METHODS

### Taxonomy

Species citation follows International Code of Zoological Nomenclature. Tardigrade taxonomy follows Bertolani et al. (2014) and later updates for Isohypsibiidae (Gąsiorek et al., 2019).

### Chemicals

Chemicals applied in functional analysis of AOX and anhydrobiosis protocol: BHAM (benzohydroxamic acid; #412260), MitoTEMPO ((2-(2,2,6,6-Tetramethylpiperidin-1-oxyl-4-ylamino)-2-oxoethyl)triphenylphosphonium chloride; #SML0737) and KCN (potassium cyanide; #60178) from Sigma-Aldrich; chemicals applied in *Saccharomyces cerevisiae* Meyen ex E.C. Hansen, 1883 cultures: Difico Yeast Extract (BD Biosciences; #212750), Bacto Peptone (BD Biosciences; #211677), D-glucose (POCH S.A. #200-075-1), glycerol (Sigma-Aldrich; #G2025) galactose (Sigma-Aldrich; #G0625), Difico Yeast Nitrogen Base without Amino Acids (BD Biosciences; #231810), Yeast Synthetic Drop-out Medium Supplements without Uracil (Sigma-Aldrich; #Y1501); chemicals for PCR and PCR product purification: Phusion® High-Fidelity PCR Kit (#E0553S) and Monarch® PCR & DNA Cleanup Kit (#T1030S), respectively from New England Biolabs.

### Bioinformatic analysis of the M. inceptum predicted AOX encoding gene product

The FASTA file of the contig containing the *M. inceptum AOX* gene and the corresponding GFF file containing the genomic location of the gene, transcript, exons and coding DNA sequences (CDS) were a kind gift of Dr Felix Bemm (Max Planck Institute for Developmental Biology, Tübingen, Germany). The obtained nucleotide sequence was translated into corresponding amino acid sequence by EMBOSS Transeq (www.ebi.ac.uk/Tools/st/emboss_transeq/) that was used to search for motives reported for animal AOX (e.g. McDonald et al., 2009; Pennisi et al., 2016) by MotifFinder being a part of I-TASSER based analysis ((Iterative Threading ASSEmbly Refinement) method (Roy et al., 2010; Yang et al., 2015; Yand and Zhang, 2015) and cell localization by DeepLoc-1.0 (www.cbs.dtu.dk/services/DeepLoc/index.php). Then the predicted amino acid sequence was aligned with AOX sequences stored in GenBank and other databases using Clustal Omega (Sievers et al., 2011) and the resulted alignment was visualized by ESPript 3.0 (Robert and Gouet, 2014). The aligned sequences were as follows: *Ciona intestinalis* (Linnaeus, 1767), (TIGR genome (Dehal et al., 2002)), *Nematostella vectensis* Stephenson, 1935 (NCBI; XM 001635879), *Lingula anatine* Lamarck, 1801 (NCBI; XP_013379624.1*), Branchiostoma floridae* Hubbs 1922 (JGI genome (Putnam et al., 2008)), *Crassostrea gigas* (Thunberg, 1793) (NCBI; FJ607013), *Dictyostelium discoideum* Raper, 1935 (NCBI; BAB82989), *Nicotiana tabacum* Linnaeus, 1753 (NCBI; AAC60576), *Zea mays* Linnaeus, 1753 (NCBI; NP001352679), *Neurospor*a *crassa* Shear and Dodge, 1927 (AAC37481) and *Trypanosoma brucei brucei* Plimmer and Bradford, 1899 (NCBI; AAB46424). To build the corresponding structural model of the predicted *M. inceptum* AOX I-TASSER was applied. The model was superimposed on the three-dimensional structure of the *T. brucei brucei* (termed here for simplicity *T. brucei*) AOX (PDB code 3VVA; Shiba et al., 2013) by application of RaptorX Structure Alignment Server (Wang et al., 2013). The predicted solution was visualized using YASARA (www.yasara.org).

### Heterologous expression of M. inceptum AOX in S. cerevisiae cells

The one CRISPR/Cas9 single-guide RNA (sgRNA) targeting *GAL1* was designed to minimize off-targets by using CRISPOR online tool (http://crispor.tefor.net/). Two oligonucleotides: forward primer (5′-GTTTTAGAGCTAGAAATAGCAAGTTAAAATAAGGC-3′) and reverse primer (5′-TTGGACGGTTCTTATGTCACGATCATTTATCTTTCACTGC-3′) carrying the 20mer guide sequence of *GAL1* were used to PCR-amplify the plasmid pML104 (Laughery et al., 2015) using the Q5 site-directed mutagenesis protocol (New England Biolabs) as described by (Hu et al., 2018). The resulting plasmid pML104-GAL1 was introduced into competent *Escherichia coli* Migula, 1885 (C2987 strain) cells by heat shock transformation. The plasmid DNA was extracted from the colonies and sequenced by T3 primer.

The full-length AOX codon-optimized open reading frame with flanking *GAL1* intergenic sequences was synthesized by Biomatik (Ontario, Canada). The synthetic sequences were received in pBluescript II SK(+) cloning vector which served as a template for repair DNA amplification by PCR. For the amplification, 20 ng of template vector was mixed with 0.5 µM forward primer (5’- ACGAATCAAATTAACAACCATAGGA-3′) and reverse primer (5’- ATGTCAAGAATAGGTATCCAAAACG-3′), 200 µM dNTPs, 1 x concentrated Phusion HF buffer and 1 U Phusion DNA Polymerase (New England Biolabs). The PCR reaction was performed as follows: an initial denaturation step at 98 °C for 30 s, followed by 30 cycles of denaturation at 98 °C for 8 s, annealing at 55 °C for 10 s, and extension at 72 °C for 40 s; and a final extension step at 72 °C for 10 min. The presence of the PCR products was confirmed by gel electrophoresis and the products were then purified by Monarch® PCR & DNA Cleanup Kit (New England Biolabs).

The wild-type *S. cerevisiae* BY4741 strain (*MAT*a, *his3*Δ, *leu2*Δ, *met15*Δ, *ura3*Δ) from EUROSCARF were grown in YPD medium containing fermentable carbon source (1% yeast extract, 2% peptone, 2% glucose) medium to OD_550_ = 4. The plasmid pML104-GAL1 and repair DNA were co-transformed into yeast by electroporation under Gene Pulser Xcell (Bio-Rad) conditions: 25 µF, 200 Ω, 1.5 kV and 0.2 cm cuvette. Then the yeast cells were selected on plates containing synthetic dextrose (SD) medium containing 0.67% yeast nitrogen base without amino acids, 2% glucose and 0.12% of drop out -ura (mixture of all amino acids and purines without uracil). Genomic DNA was then extracted from the resulting colonies and the presence of AOX encoding sequence was verified by sequencing (see also S1 Fig.).

### Detection of M. inceptum AOX activity in mitochondria of S. cerevisiae intact cells

Cells of BY4741 + AOX yeast strain (*MAT*a, *his3*Δ, *leu2*Δ, *met15*Δ, *ura3*Δ, *gal1*Δ::*Mi-AOX*) were grown in YPG medium containing non-fermentable carbon source (1% yeast extract, 2% peptone, 3% glycerol at pH = 5,5) to OD_550_ = 1, then galactose at a final concentration of 2% was added to induce and maintain AOX expression for 24 h. In parallel, control yeast cells were cultured in the absence of galactose. Then, yeast cells were washed twice with distilled water, resuspended in 100 µL YPG medium and quantified by OD_550_ measurement. For the same amount of cells rates of oxygen uptake were determined in 1 mL of YPG medium using a Clark electrode (Hansatech Instruments). To determine bioenergetic parameters related to AOX activity, 1mM KCN and 3 mM BHAM were applied to inhibit the respiratory chain complex IV (cytochrome c oxidase) and AOX, respectively. The applied final concentrations of KCN and BHAM were verified experimentally to obtain saturated effect.

### Detection of M. inceptum AOX presence in S. cerevisiae isolated mitochondria

Mitochondria of *S. cerevisiae* cells expressing AOX were isolated according to the published procedure (Daum et al., 1982). Protein concentration was measured by the method of Bradford and albumin bovine serum (BSA), essentially fatty acid free, was used as a standard. The mitochondrial proteins were separated by SDS-PAGE in the presence of 6 M urea and gels were stained with Coomassie Brilliant Blue G-250.

### Culture of *M. inceptum*

The initial specimens of *M. inceptum* were collected in a xerothermic habitat, i.e. mosses on concrete wall in the city centre of Poznań in Poland (Heliodor Święcicki Clinical Hospital of Poznań University of Medical Sciences at Przybyszewskiego street, 52°24’15”N, 16°53’18”E; 87 m asl). The extraction was performed by the standard method (Dastych, 1980). To maintain the culture *M. inceptum* specimens were kept at 18 °C in the darkness and at relative humidity (RH) of 40% (POL EKO KK 115 TOP+ climatic chamber) in covered Petri dishes (5.5 cm in diameter) with bottom scratched by sandpaper to allow tardigrade movement. The animals were coated by a thin layer of the culture medium, i.e. spring water (Żywiec Zdrój; Żywiec Zdrój S.A., Poland) mixed with double distilled water (ddH_2_O) in 1:3 ratio. The culture medium was exchanged every week and the animals were fed with the nematode *Caenorhabditis elegans* (Maupas, 1900), and the rotifer *Lecane inermis* (Bryce, 1892), 1.A2.15 strain. The nematode wild type Bristol N2 strain was obtained from the Caenorhabditis Genetics Center (CGC) at the University of Minnesota (Duluth, Minnesota, USA) and its culture was maintained by standard methods (Brenner, 1974).

### Anhydrobiosis protocol

Groups of 10 fully active, (displaying coordinated movements of the body and legs) adult specimens of medium body length of about 500-550 µm (in 4-6 replicates), cleaned of debris, were transferred and dehydrated in 400 µl of the culture medium in 3.5 cm (in diameter) covered Petri-dishes with bottom scratched by sandpaper. The dishes were allowed to dry slowly in the Q-Cell incubator (40-50% RH, 20 °C, complete darkness) for 72h. The dehydrated animals (tuns) were kept under the above conditions for 3, 30 or 60 days. Tun formation and the control and rehydrated specimen activity were monitored under Olympus SZ61 stereomicroscope connected to Olympus UC30 microscope digital camera. The obtained images and films were transferred to a computer and processed by CellSens Standard (Olympus) software.

After the mentioned duration of the tun stage, 3 ml of the culture medium were added to each dish with tuns and the tuns were transferred to small glass cubes in which they were observed under stereomicroscope for 24h, and the animal return to full activity was monitored. The average number of specimens able to recover to full activity after 24h from the onset of rehydration was defined as final return to full activity.

When indicated, BHAM (prepared in methanol) and MitoTEMPO (prepared in ddH_2_O) were added at the beginning of animal dehydration or rehydration stages. The applied final concentrations of BHAM (0.1 and 0.2 mM) and MitoTEMPO (0.01 mM) were verified experimentally to avoid lethal effect. Because two different solvents were applied for the studied compounds, two different controls were used in experiments, namely control 1 (C_1_) being suitably diluted methanol solution (0.3% v/v) and control 2 (C_2_) being clean ddH_2_O.

### Statistical analysis

Effects of BHAM and KCN on the rate of oxygen uptake, displayed by *S. cerevisiae* cells was analyzed by t-test. For testing differences in number of tuns formed under different conditions (in the presence of MitoTEMPO or BHAM as well as for proper controls) one-way ANOVA (analysis of variance) (Zar, 1999) was applied. To test differences in animal return to full activity between two grouping variables: (I) 12 *‘time windows*’ (10, 20, 30, 40, 60, 90, 120, 180, 240, 360, 480, 1440 min) and (II) three ‘*experimental groups*’ (control 1 (C_1_), 0.1 mM BHAM and 0.2 mM BHAM) or (III) two “*experimental groups*” (control 2 (C_2_) and 0.01 mM MitoTEMPO)), Factorial ANOVA was applied (Zar, 1999). We used the standard presentation of results for this type of analysis (Zar, 1999; Sokal and Rohlp, 1995). For each model we calculated *F*-statistics and R^2^ as a measure of variance explained (Sokal and Rohlp, 1995). For ANOVA models, when *F* was statistically significant, in the next step we deepened that analysis by calculation of t-test for each grouping variables. If the *t*-test for the factor ‘*experimental groups*’ was statistically significant, it provides (from a methodical point of you) an authorization to perform *Tukey post-hoc test* (Zar, 1999) to test the obtained differences. Finally post-hoc test results were visualized in figures. To compare effects imposed by BHAM and MitoTEMPO juxtaposed to proper controls on anhydrobiotic animal return to full activity, regardless of the “*time windows*”, we developed Linerar Mixed Models for applied duration of the tun stage (3 and 30 days) and applied concentrations of BHAM (0.1 mM and 0.2 mM) and MitoTEMPO (0.01 mM). We used *concentration* as fixed effect whereas for random effect we used *time window*. All calculations were performed by R (R Core Team 2013).

## RESULTS

### The predicted M. inceptum AOX is a functional protein

The predicted *M. inceptum* AOX amino acid sequence revealed two properties characteristic for animal AOX (Fig. 1A and S1 Data). They were: (i) the presence of conserved glutamate (E) and histidine (H) residues (marked with stars) within the ferritin-like domain (blue box), required for two iron atom binding and crucial for functionality of the catalytic core and (ii) the presence of N-P-[YF]-X-P-G-[KQE] motif at C-terminus (violet box) which is regarded as the diagnostic for animal AOX identification (McDonald et al., 2009; Pennisi et al., 2016) although the last variable position was occupied by arginine (R).

**Fig. 1.**
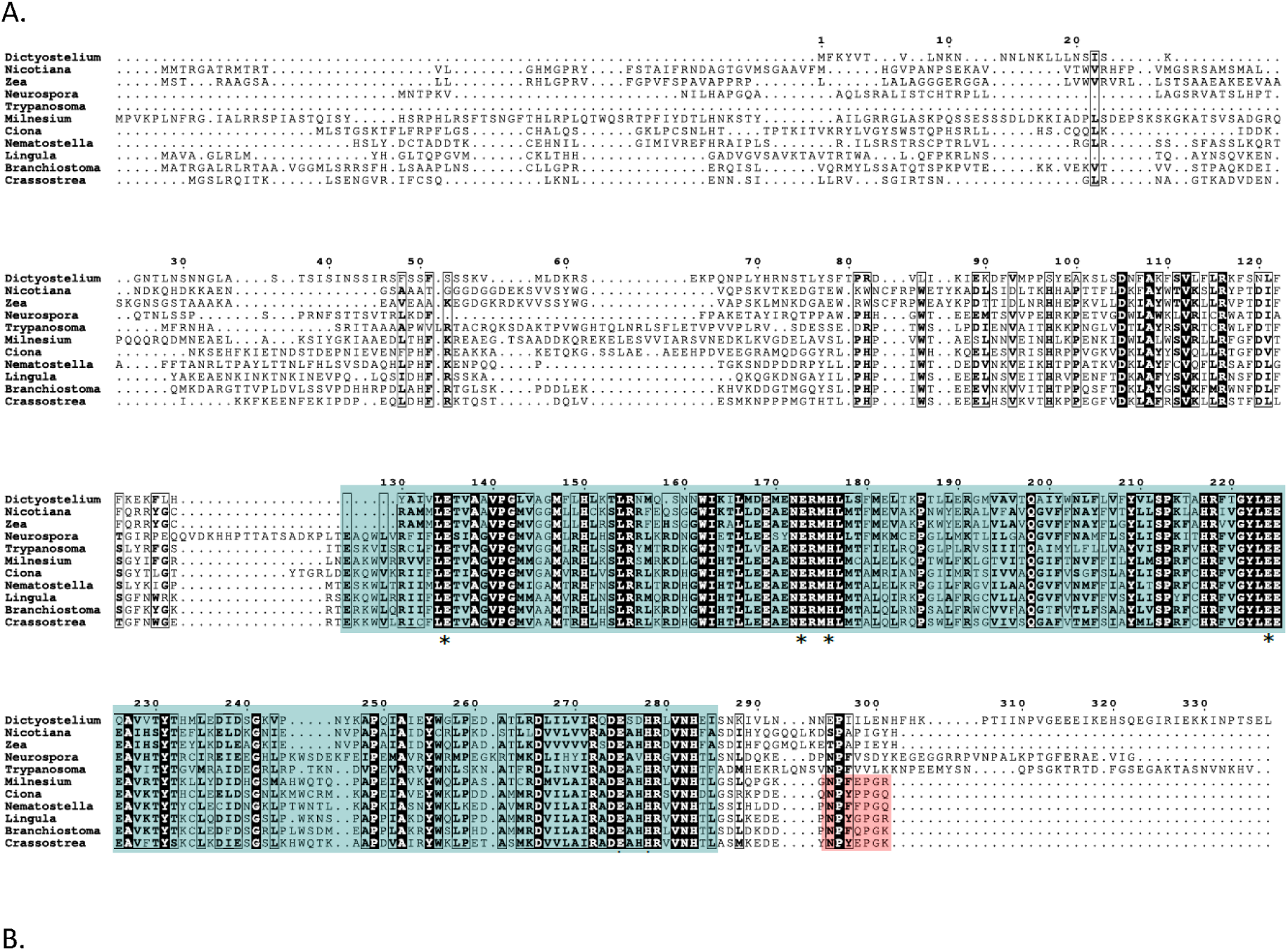
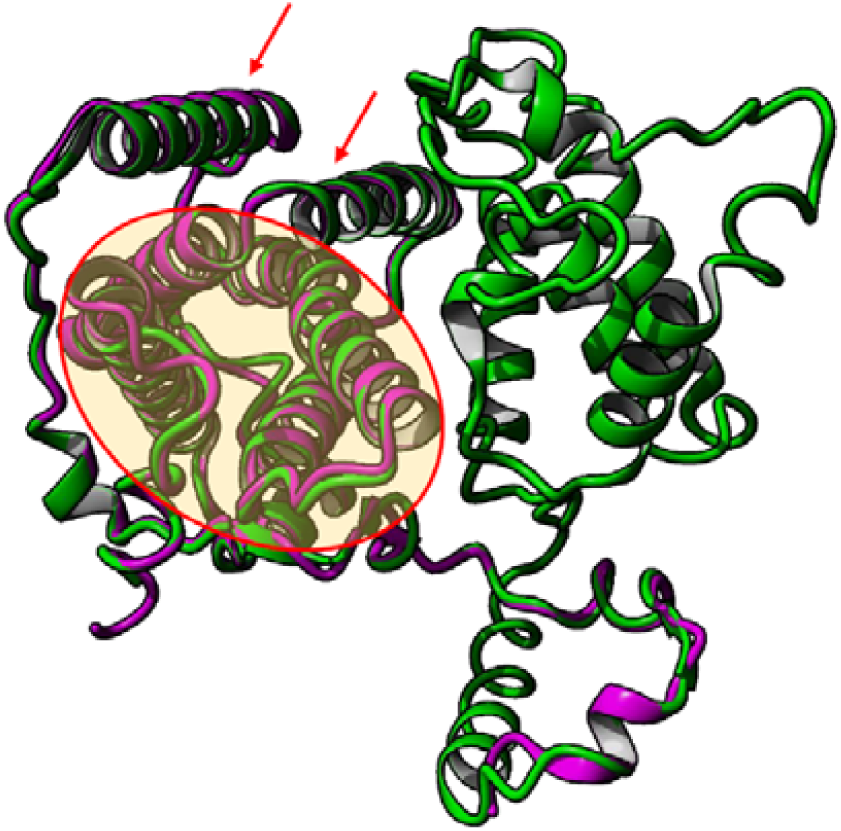
Bioinformatic identification of *Milnesium inceptum* AOX. (A) A multiple sequence alignment of full-length alternative oxidase (AOX) proteins from selected animal species and representatives of other phylogenetic lineages. The conserved glutamate (E) and histidine (H) residues within the ferritin-like domain (blue box) are marked with stars. The presence of N-P-[YF]-X-P-G-[KQE] motif at C-terminus regarded as diagnostic for animal AOX identification is marked by violet box. Animal species (Opisthokonta) are represented by *Ciona intestinalis, Nematostella vectensis, Lingula anatina, Branchiostoma floridae, and Crassostrea gigas*. The other phylogenetic lineages are represented by *Dictyostelium discoideum* (Amoebozoa), *Nicotiana tabacum* and *Zea mays* (Archaeplastida, Plants), *Neurospora crassa* (Opisthokonta, Fungi) and *Trypanosoma brucei brucei* (termed here for simplicity *T. brucei*) (Excavata). The species taxonomy follows Keeling et al., 2005. (B) Comparison of 3D structure resolved for *T. brucei* AOX (3VVA) and predicted for *M. inceptum* AOX. Red frame and arrows indicate four-α-helix bundle of the catalytic core and two additional α-helices anchoring the protein to the membrane, respectively. purple, *T. brucei brucei* AOX monomer; green, *M. inceptum* protein.

To verify the conservation of the predicted *M. inceptum* AOX amino acid sequence three-dimensional structure modeling of the protein was performed and the result compared with the *T. brucei* AOX three-dimensional structure (PDB code 3VVA) (Shiba et al. 2013). As shown in Fig. 1B (see also S1 Data), the *M. inceptum* amino acid sequence (green) was able to form six α-helices. They appeared to match up the corresponding regions in the structural model of *T. brucei* AOX (blue), i.e. four-α-helix bundle of the catalytic core, hosting the di-iron catalytic site, and to two additional α-helices flanking the catalytic core and anchoring the protein to the membrane (Pennisi et al., 2016). Accordingly, DeepLoc-1.0 based analysis (S1 Data) indicated the presence of N-terminal sequence enabling the predicted protein to be localized in the mitochondrial inner membrane (Tanudji et al., 1999).

To functionally access *M. inceptum* AOX, the predicted protein was expressed in the yeast *S. cerevisae* cells under the *GAL1* promoter control. The cells were grown in YPG medium containing non-fermentable carbon source (glycerol) metabolized by respiration that requires functioning mitochondria (Gałganska et al., 2008). The expression of *M. inceptum* AOX was induced and maintained within 24 h by addition of galactose. Then cell respiration, i.e. the rate of oxygen uptake displayed by the cells was estimated and the effects of known inhibitor of AOX (i.e. BHAM) and the cytochrome c oxidase representing the respiratory chain cytochrome pathway (i.e. KCN) were determined. In the absence of *M. inceptum* AOX expression (Fig. 2A), the yeast cell respiration was totally inhibited by 1 mM KCN but not by 3 mM BHAM, independently of the order of their additions. In the presence of *M. inceptum* AOX expression (Fig. 2B), the inhibitory effect of KCN was distinctly diminished and the resulting KCN-resistant respiration was sensitive to BHAM. Importantly, BHAM-sensitive respiration was also observed before KCN addition. As summarized in Fig. 2C, respiration of *S. cerevisiae* cells not expressing *M. inceptum* AOX was sensitive to KCN but not to BHAM whereas respiration of the yeast cells expressing *M. inceptum* AOX was partially sensitive to KCN and was also sensitive to BHAM. Moreover, basal respiration of *S. cerevisiae* cells expressing *M. inceptum* AOX was approximately two times higher but the difference was eliminated by addition of BHAM. To verify correct targeting of *M. inceptum* AOX to *S. cerevisiae* mitochondria, the isolated mitochondria were separated by SDS-PAGE and analysed by the standard gel staining. As shown in Fig. 2D, mitochondria isolated from the yeast cell cultured in the presence of galactose triggering the AOX expression contained distinct amounts of a protein of molecular weight close to 40 kD which corresponds to heterologously expressed animal AOX proteins (Robertson et al., 2016).

**Fig. 2.**
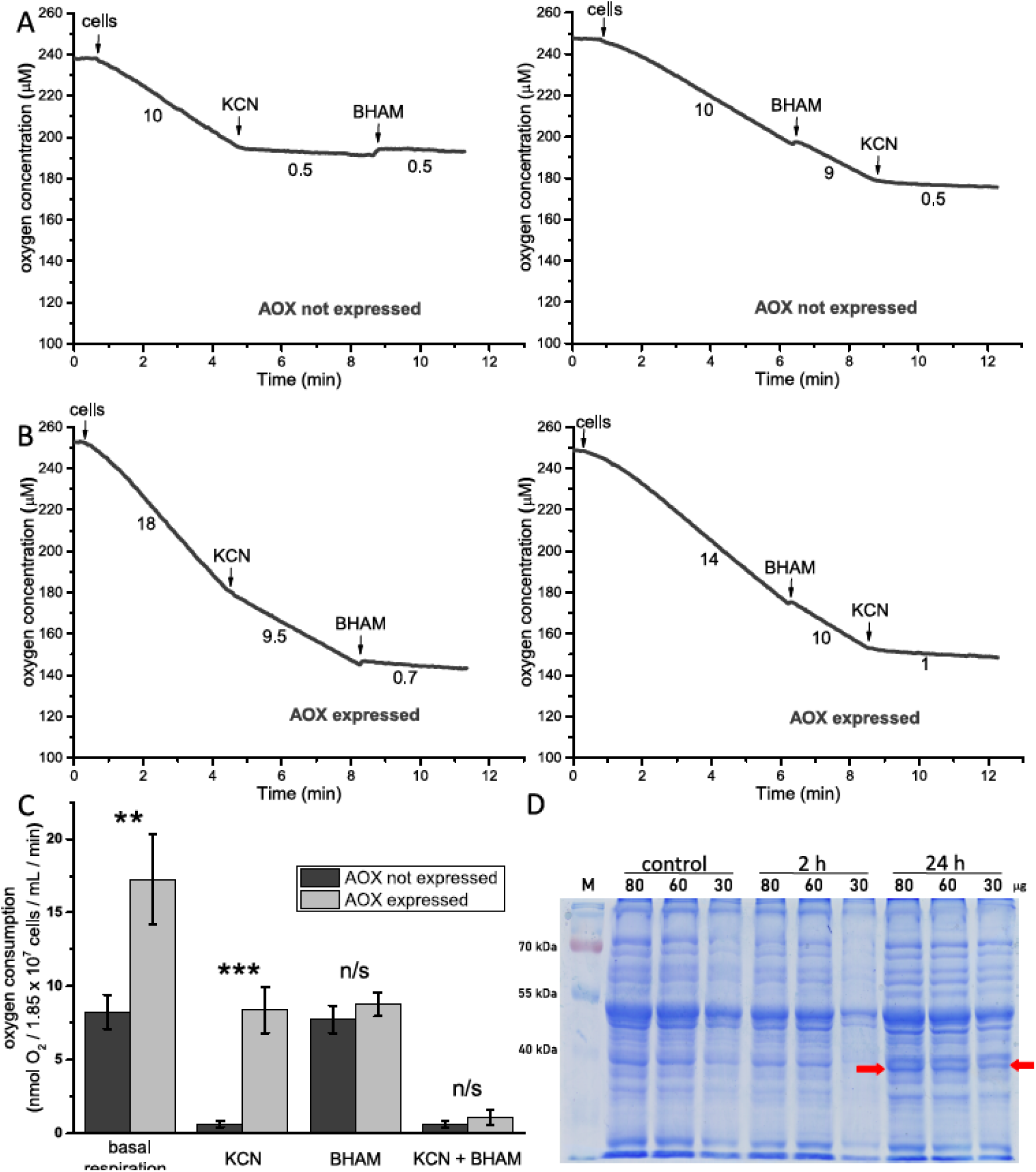
Functional analysis of *Milnesium inceptum* AOX expressed in *Saccharomyces cerevisiae* mitochondria. The tardigrade AOX expression was induced by the addition of galactose to YPG medium. (A) and (B) Representative traces of the performed measurements of the rate of oxygen uptake by intact yeast cells in the absence (AOX not expressed) and in the presence (AOX expressed) of galactose. (C) Changes in *S. cerevisiae* cell respiration in the presence or the absence of AOX expression and after addition of inhibitors of AOX (BHAM) and the respiratory chain complex IV (KCN). The applied concentrations of inhibitors were as follows: 1mM KCN and 3 mM BHAM. ** *p* ⍰ 0.01; *** *p* ⍰ 0.001; n/s not statistically significant. (D) Detection of *M. inceptum* AOX expression and maintenance in *S. cerevisiae* mitochondria in the presence of galactose. Mitochondria were isolated from cells cultured in the absence of galactose (control) and in its presence, after 2 and 24 h. The red arrows indicate bands corresponding to *M. inceptum* AOX.

### The appearance of M. inceptum tuns is not changed by their formation in the presence of BHAM and MitoTEMPO

To validate putative contribution of AOX to anhydrobiosis we first checked the BHAM presence effect on the formed tun appearance. Because hydroxamic acids, including BHAM, are also known to display free-radical scavenging activity (e.g. Przychodzen et al., 2013; Adewuyi et al., 2015), we compared the BHAM effect with that imposed by the presence of MitoTEMPO, a well-known mitochondria-specific superoxide scavenger (Fig. 3). The expected appearance of tuns was compact body shape resulting from contracting the body and withdrawal of the legs into the body cavity accompanied by the body water loss (Møbjerg et al., 2018). The presence of 0.1 or 0.2 mM BHAM and 0.01 mM MitoTEMPO as well as the proper dilution of methanol used as BHAM solvent (control 1, C_1_) did not cause statistically significant differences in the average number of tuns with the expected appearance when compared with control 2 (C_2_) being clean water. Thus, the presence of the applied concentrations of BHAM and MitoTEMPO did not affect tun appearance.

**Fig. 3.**
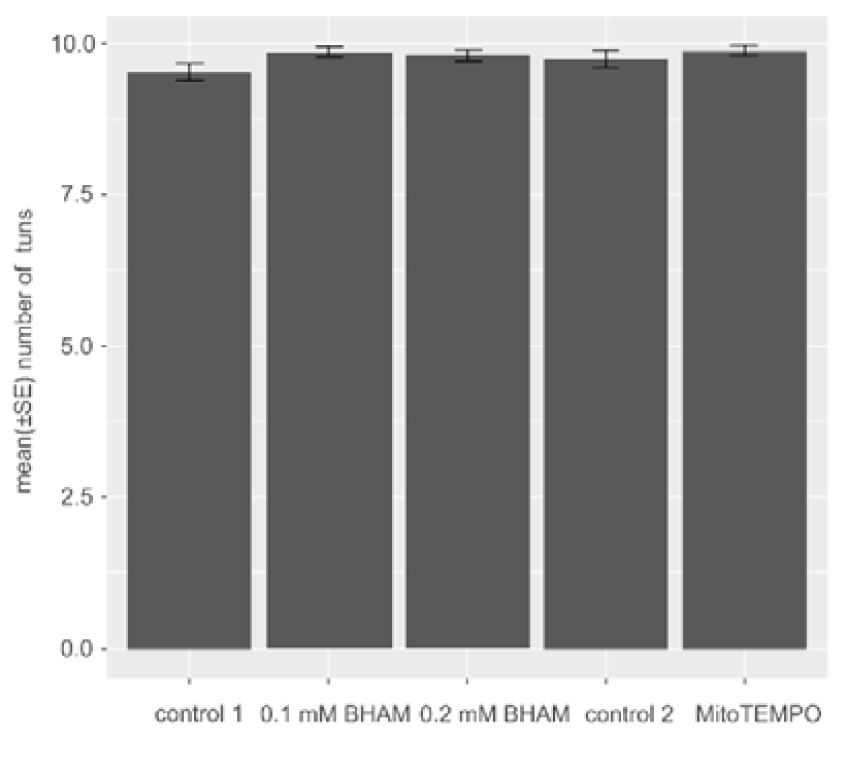
The average numbers of tuns formed by *M. inceptum* specimens in the absence and in the presence of BHAM and MitoTEMPO. To obtain the tun stage groups of 10 fully active adult specimens were dehydrated in the absence and in the presence of BHAM (0.1 or 0.2 mM) and MitoTEMPO (0.01 mM). The full activity was defined as coordinated movements of animal body and legs (crawling). control 1, 0.3% methanol; control 2, ddH_2_O. The differences between the means are not statistically significant (one-way ANOVA: F_4,89_ = 1.65 *p* = 0.167).

### The presence of BHAM during tun formation but not during tun rehydration affects M. inceptum specimen recovery to full activity

It is well known that the longer a specimen is in the tun stage, the longer period it requires to return to active life (e.g. Wright, 1989; Rebecchi et al., 2007; Schill and Hengherr, 2018). The same was observed for the studied *M. inceptum* specimens (Fig. 4). As expected for the applied duration of the tun stage, i.e. 3, 30 and 60 days, the revival corresponding to return to full activity including the final return (defined as the average number of specimens able to recover to full activity after 24h from the onset of rehydration) was delayed as the tun stage duration extended. The effect of BHAM presence added during tun formation or during tun rehydration on anhydrobioic animal recovery to full activity for different duration of the tun stage was analysed using Factorial ANOVA (Tab. 1). It should be mentioned that statistical analysis of the performed experiments assumed two grouping variables, i.e. time windows and two different concentrations of BHAM versus control conditions. Thus, it was impossible to simply test the differences between each group as the effect of “multiple comparisons” would decrease the biological value of the results (Sokal and Rohlp, 1995).

**Fig. 4.**
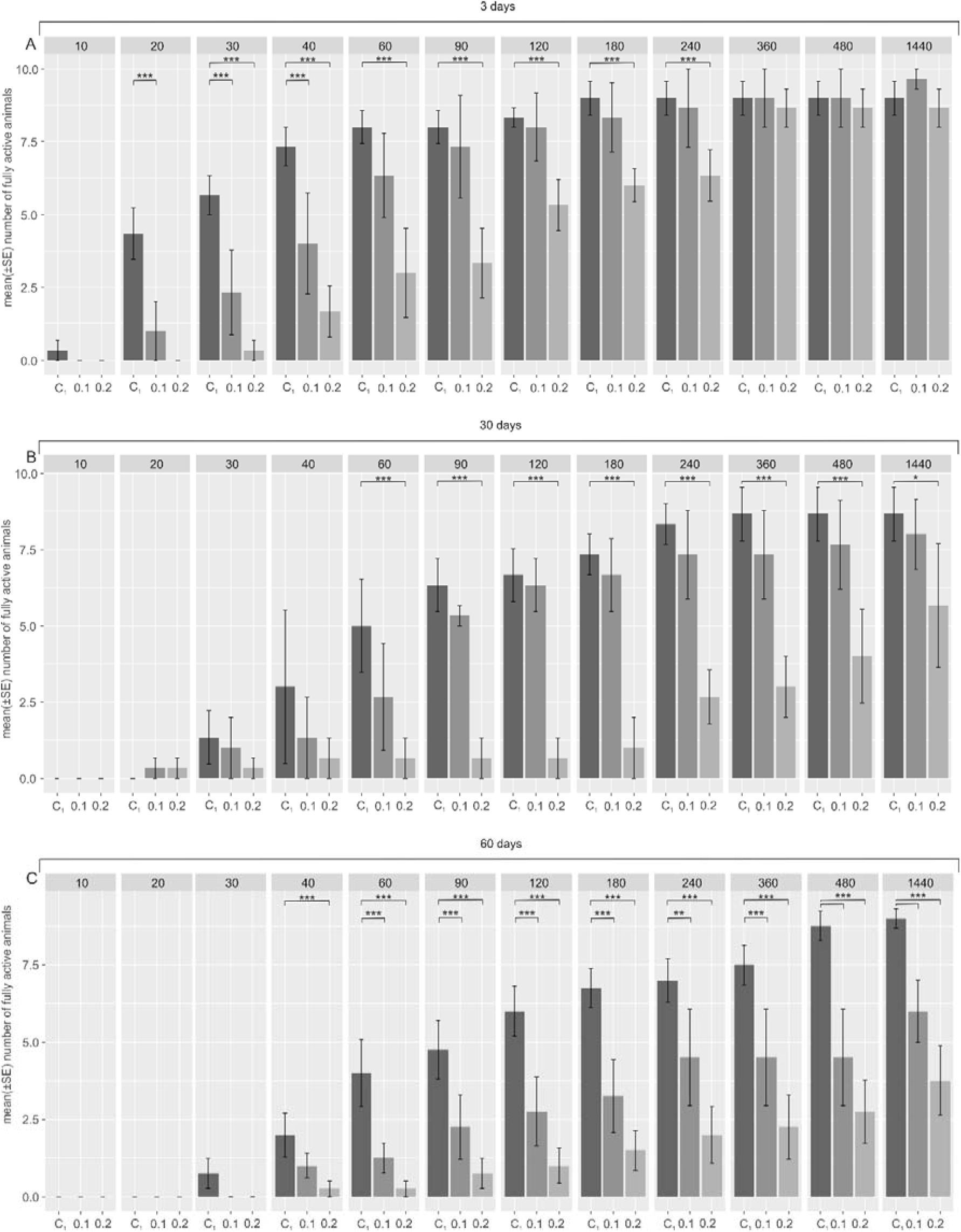
The effect of the BHAM presence during *Milnesium inceptum* tun formation on the specimen return to full activity. (A), (B) and (C) Tun duration for 3, 30 and 60 days, respectively. Full activity was defined as coordinated movements of animal body and legs (crawling). The time points of observations are indicated in minutes following the onset of rehydration. C_1_, control 1 (0.3% methanol); 0.1 and 0.2, two different BHAM concentrations, i.e. 0.1 and 0.2 mM BHAM, respectively. * *p* ⍰ 0.05; ** *p* ⍰ 0.01; *** *p* ⍰ 0.001 (see also Tab. 1).

**Tab. 1.** Results of Factorial ANOVA. For each experiment the models included two factors, i.e. (I) 12 time windows (10, 20, 30, 40, 60, 90, 120, 180, 240, 360, 480, 1440 min) and (II) three experimental groups (control, 1 mM BHAM and 0.2 mM BHAM). *F-test*, variation between sample means/variation within the samples; DF, mean degree of freedom used to variance calculations; *p*, level of α significance; R^2^, variance explained; *t*, statistics for factor inside ANOVA model (see also Fig. 4-5).

| model | Factorial ANOVA model |  |  | Factors in models |  |  |
| --- | --- | --- | --- | --- | --- | --- |
|  | *F*-test<sub>(DF)</sub> | *p* | *R*<sup>2</sup> | Factor | *t* | *p* |
| Dehydration<br>3 day tuns | 28.21<sub>(2,105)</sub> | <0.001 | 0.33 | Time window | 6.15 | < 0.001 |
|  |  |  |  | Experimental group | -4.30 | <0.001 |
| Dehydration<br>30 day tuns | 39.78<sub>(2,105)</sub> | <0.001 | 0.42 | Time window | 6.75 | <0.001 |
|  |  |  |  | Experimental group | -5.82 | <0.001 |
| Dehydration<br>60 day tuns | 67.92<sub>(2,141)</sub> | <0.001 | 0.48 | Time window | 8.60 | <0.001 |
|  |  |  |  | Experimental group | -7.85 | <0.001 |
| Rehydration<br>3 day tuns | 18.07<sub>(2,201)</sub> | <0.001 | 0.14 | Time window | 5.85 | <0.001 |
|  |  |  |  | Experimental group | -1.34 | 0.17 |
| Rehydration<br>60 day tuns | 23.37<sub>(2,141)</sub> | <0.001 | 0.23 | Time window | 6.67 | <0.001 |
|  |  |  |  | Experimental group | 1.48 | 0.14 |

Because in the case of the BHAM presence during tun formation the *t*-test for the factor ‘*experimental group*’ (i.e. control 1 (C_1_), 1 mM BHAM and 0.2 mM BHAM) was statistically significant we developed the Tukey post-hoc test (Zar, 1999) to test differences between (I) C_1_ *vs*. 0.1 mM BHAM and (II) C_1_ *vs*. 0.2 mM BHAM (Tab. 1). As shown in Fig. 4, the presence of 0.1 or 0.2 mM BHAM, during tun formation additionally affected the return to full activity in a way dependent on BHAM concentration. For the tun stage lasting 3 days (Fig. 4A) statistically significant distinct delay was observed in return to full activity, especially for higher BHAM concentration, although the final return to full activity was comparable to the specimens from the control group. However, the final return to full activity was distinctly decreased for the tun stage formed in the presence of BHAM and lasting 30 and 60 days. In the case of the former (30 days) the clear statistically significant decrease was observed only for 0.2 mM BHAM (Fig. 4B) whereas in the case of the latter (60 days) for both BHAM concentrations and the decrease was more pronounced for 0.2 mM BHAM (Fig.4C). Thus, duration of the tun stage as well as the presence of BHAM during tun formation and consequently during the tun stage imposed statistically significant additive effect on anhydrobiotic animal return to full activity which also depended on the AOX inhibitor concentration.

The same approach of data analysis applied for the presence of BHAM during tun rehydration resulted in the *t*-test for factor ‘*experimental groups*’ (i.e. (C_1_), 1 mM BHAM and 0.2 mM BHAM) that was not statistically significant (Tab. 1 and Fig. 5). It denoted that when added during rehydration BHAM did not influence return to full activity in a statistically significant way and that concerned different tun stage duration. Thus, the presence of BHAM during tun rehydration appeared to not affect animal return to full activity.

**Fig. 5.**
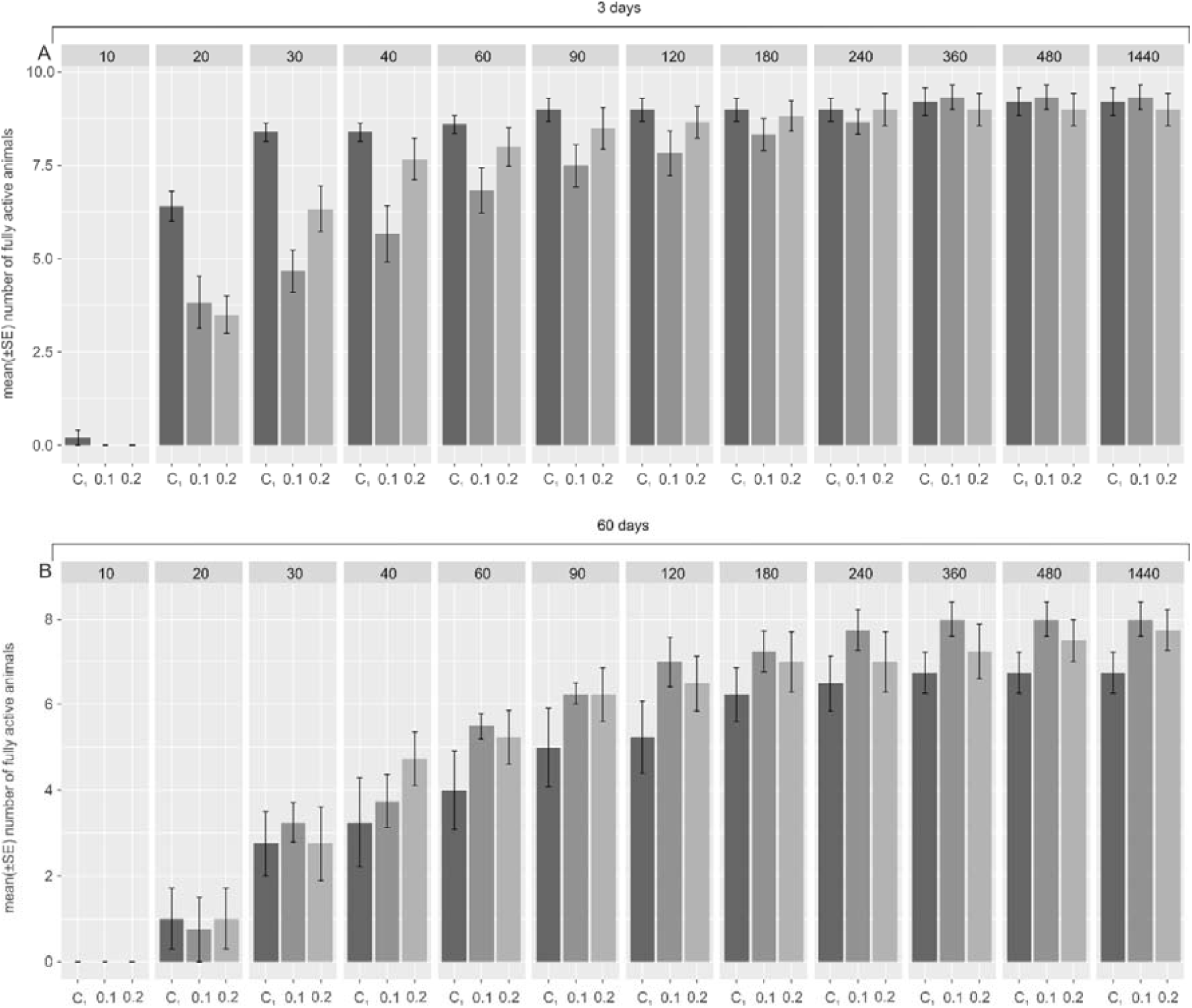
The effect of the BHAM presence during *Milnesium inceptum* tun rehydration on the specimen return to full activity. (A) and (B) Tun duration for 3 and 60 days, respectively. Full activity was defined as coordinated movements of animal body and legs (crawling). The time points of observations are indicated in minutes following the onset of rehydration. C_1_, control 1 (0.3% methanol); 0.1 and 0.2, two different BHAM concentrations, i.e. 0.1 and 0.2 mM BHAM, respectively. The differences observed for the mentioned experimental groups were not statistically significant (see also Tab. 1).

### BHAM and MitoTEMPO applied during tun formation display different effects on M. Inceptom specimen recovery to full activity

Since data shown in Fig. 4 indicated that the presence of BHAM during tun formation affected anhydrobiotic animal return to full activity, we decided to check free-radical scavenging activity of BHAM. For that purpose we analyzed the effect of 0.01 mM MitoTEMPO, a well-known mitochondria-specific superoxide scavenger, added during tun formation on anhydrobiotic tardigrade recovery to full activity for different duration of the tun stage (3 and 60 days) using Factorial ANOVA (Tab. 2). As illustrated in Fig. 6, the presence of MitoTEMPO during tun formation resulted in the *t*-test for factor ‘*experimental groups*’ (i.e. control 2 (C_2_) and 0.01 mM MitoTEMPO) that was not statistically significant.

**Tab. 2.** Results of Factorial ANOVA. For each experiment the models included two factors, i.e. (I) 12 time windows (10, 20, 30, 40, 60, 90, 120, 180, 240, 360, 480, 1440 min) and (II) two experimental groups (control, 0.01 mM MitoTEMPO). *F-test*, variation between sample means/variation within the samples; DF, mean degree of freedom used to variance calculations; *p*, level of α significance; R^2^, variance explained; *t*, statistics for factor inside ANOVA model (see also Fig. 6).

| model | Factorial ANOVA model |  |  | Factors in models |  |  |
| --- | --- | --- | --- | --- | --- | --- |
|  | *F*-test<sub>(DF)</sub> | *p* | R<sup>2</sup> | Factor | *t* | *p* |
| **Dehydration**<br>**3 day tuns** | 4.55<sub>(2.57)</sub> | 0.014 | 0.10 | Time window | 2.78 | 0.007 |
|  |  |  |  | Experimental group | 1.16 | 0.250 |
| **Dehydration**<br>**60 day tuns** | 2.26<sub>(2.81)</sub> | <0.001 | 0.36 | Time window | 7.05 | <0.001 |
|  |  |  |  | Experimental group | 0.84 | 0.399 |

**Fig. 6.**
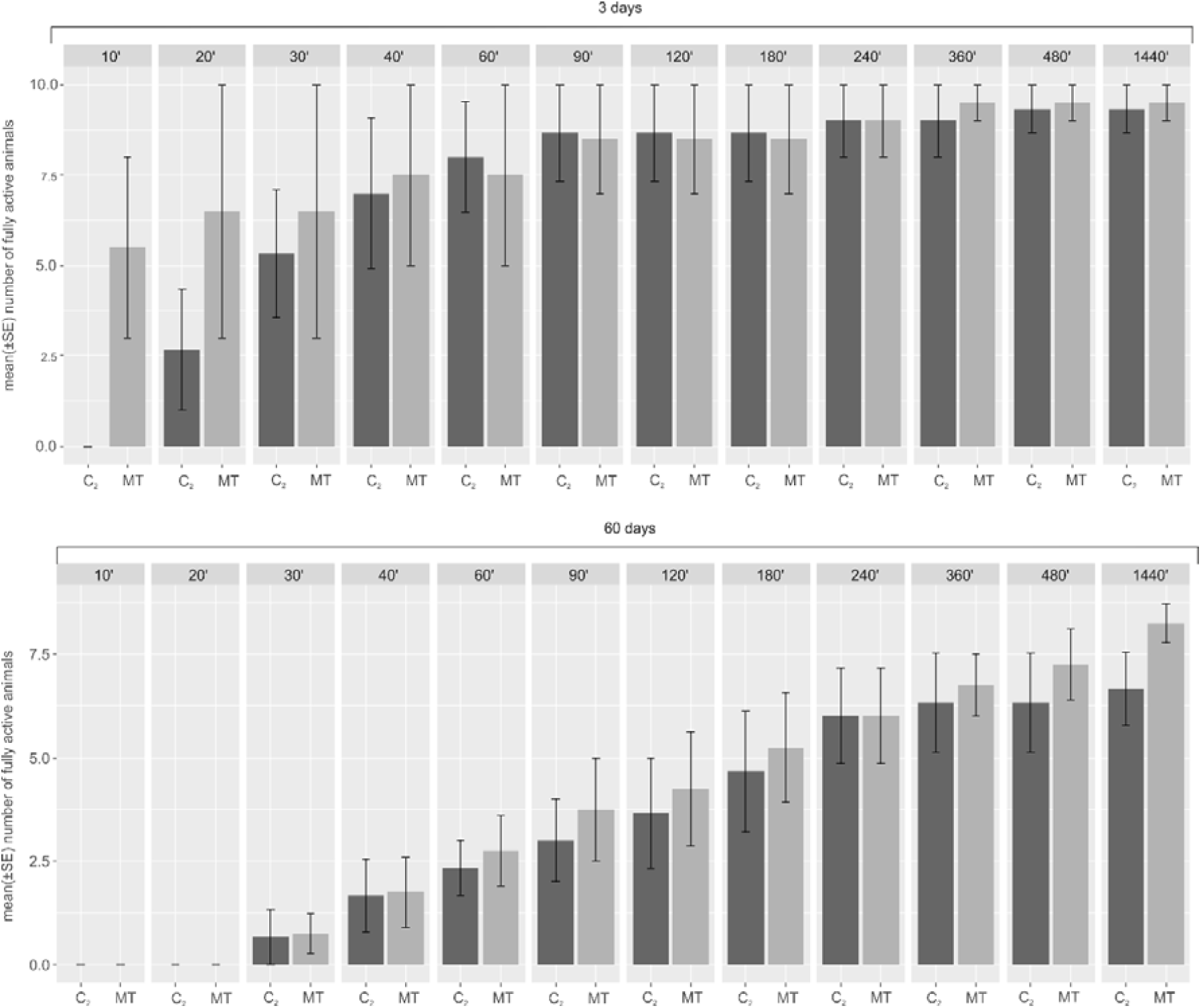
The effect of the MitoTEMPO presence during *Milnesium inceptum* tun formation on the specimen return to full activity. (A) and (B) Tun duration for 3 and 60 days, respectively. Full activity was defined as coordinated movements of animal body and legs (crawling). The time points of observations are indicated in minutes following the onset of rehydration. C_2_, control (ddH_2_O); MT, 0.01 mM MitoTEMPO. The differences observed for the mentioned experimental groups were not statistically significant (see also Tab. 2).

Then we compared the observed effects for the BHAM and MitoTEMPO treated tun forming animals. For that we developed Linear Mixed Models enabling analysis of ratios between average numbers of fully active animals obtained for BHAM or MitoTEMPO treated tardigrades and proper control animals observed in different time points of animal activity observation following rehydration. As summarized in Tab. 3 and shown in Fig. 7, for the applied duration of the tun stage (3 and 60 days), the presence of MitoTEMPO during tun formation did not affect return to full activity in a statistically significant way. Nevertheless, animals forming tuns in the presence of MitoTEMPO displayed clearly faster return to full activity that animals forming tuns in the presence of BHAM. Furthermore, the BHAM effect was distinctly dependent on its concentration and the tun stage duration. Thus, the effects of BHAM and MitoTEMPO on tardigrade recovery to full activity were different when these reagents were present during tun formation. This observation suggested that effects imposed by BHAM appeared to be based on different mechanisms than that of MitoTEMPO.

**Tab. 3.** Results for Linear Mixed Models developed to compare effects imposed by BHAM and MitoTEMPO juxtaposed to proper controls on average numbers of revived animals, regardless of the “time windows”. The models were performed for applied duration of the tun stage (3 and 30 days) and applied concentrations of BHAM (0.1 mM and 0.2 mM) and MitoTEMPO (0.01 mM). Concentration was used as fixed effect and “time window” was used for random effect. *F-test*, variation between sample means / variation within the samples; *p*, level of α significance; *t*, statistics for factor inside Linear Mixed Model (see also Fig. 7).

| Group |  | Linear Mixed Model |  | Variable inside the Linear Mixed Model |  |  |  |
| --- | --- | --- | --- | --- | --- | --- | --- |
|  |  | *F* | *p* | variables | *t* (for fixed variable) | Chi.sq (for random variable) | *p* |
| Dehydration<br>3 day tuns | 0.1 mM BHAM<br>/ 10 μM MT | 9.01 | < 0.001 | Experimental group | -3.00 | - | 0.004 |
|  |  |  |  | Time window | - | 17.86 | <0.001 |
| Dehydration<br>3 day tuns | 0.2 mM BHAM<br>/ 10 μM MT | 9.03 | 0.004 | Experimental group | -2.73 | - | <0.006 |
|  |  |  |  | Time window | - | 10.7 | 0.003 |
| Dehydration<br>60 day tuns | 0.1 mM BHAM<br>/ 10 μM MT | 7.00 | <0.001 | Experimental group | 3.02 | - | <0.001 |
|  |  |  |  | Time window | - | 12.7 | <0.001 |
| Dehydration<br>60 day tuns | 0.2 mM BHAM<br>/ 10 μM MT | 12.89 | <0.001 | Experimental group | 6.78 | - | <0.001 |
|  |  |  |  | Time window | - | 13.65 | <0.001 |

**Fig. 7.**
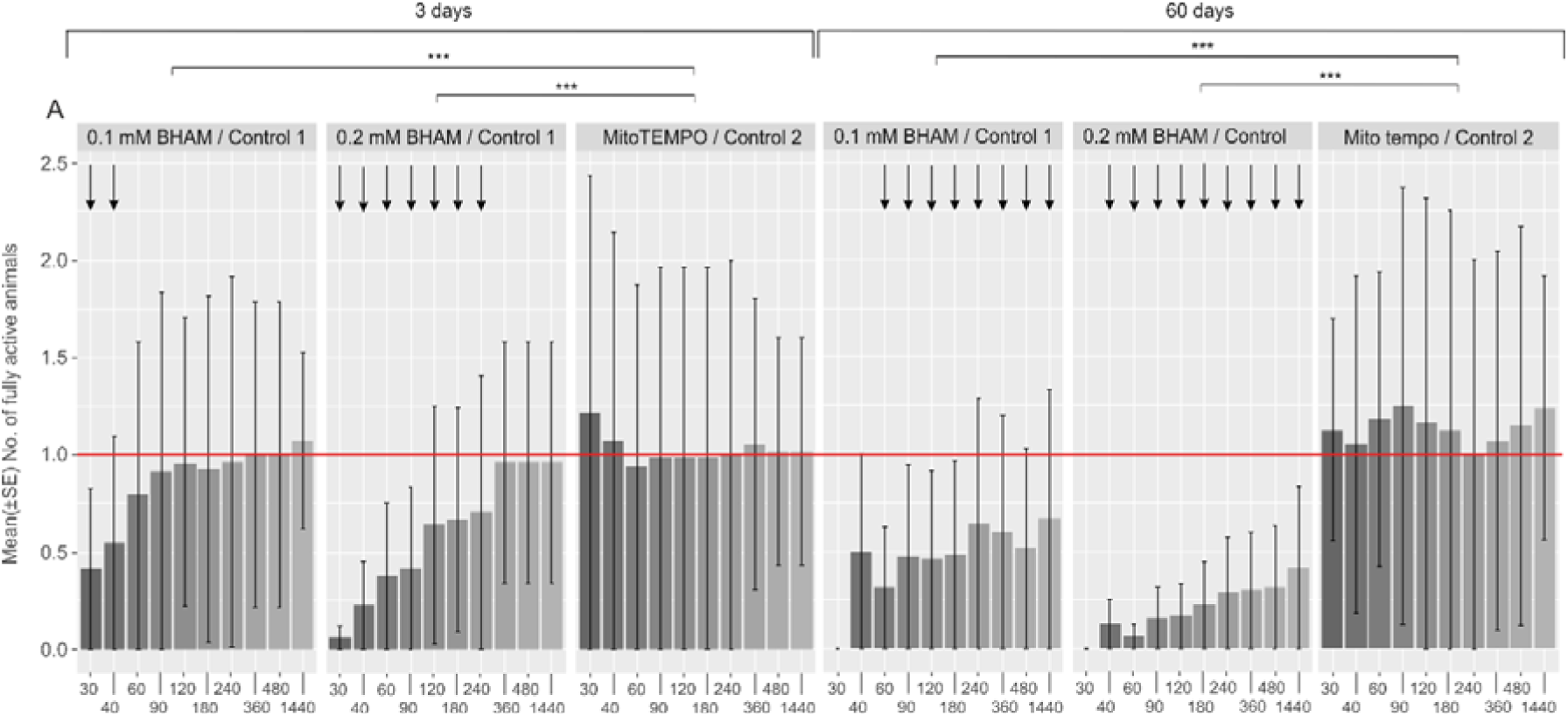
Comparison of the effect of the BHAM and MitoTEMPO presence during *Milnesium inceptum* tun formation on the specimen return to full activity. For the comparison ratios between average numbers of fully active animals obtained for BHAM (0.1 or 0.2 mM) and MitoTEMPO (0.01 mM) treated animals and proper control animals were calculated and analysed by application of the developed Linear Mixed Models (see Tab. 3). The analysis was performed for different time points of the animal observation (in minutes) following the onset of rehydration. Full activity was defined as coordinated movements of animal body and legs (crawling). control 1, 0.3% methanol; control 2, ddH_2_O. *** *p* ⍰ 0.001 (see also Tab 3). Arrows indicate statistically significant differences between the experiment groups, shown also in Fig. 4.

## DISCUSSION

Here we provide data on functionality of the tardigrade *Milnesium inceptum* AOX and its possible contribution to the animal anhydrobiosis. The presence of functional AOX in the tardigrade mitochondria was confirmed bioinformatically and experimentally. The predicted full amino acid sequence of the AOX appears to be imported into the inner mitochondrial membrane as well as contains the ferritin-like domain and characteristic motif at C-terminus defined as marker of animal AOX (Fig.1A and S1 Data). Moreover, the sequence is able to form tertiary structure postulated for AOX (Fig.1B and S1 Data) that includes four-α-helix bundle of the catalytic core, hosting the di-iron catalytic site, and two additional α-helices flanking the catalytic core (McDonald et al., 2009; Pennisi et al., 2016). After heterologous expression in *S. cerevisiae* cells, *M. inceptum* AOX confers KCN-resistant and AOX inhibitor-sensitive respiration to these cells grown on non-fermentative carbon sources (Fig.2). The analogous observation was reported for the pacific oyster *C. gigas* AOX heterologously expressed in the yeast mitochondria (Robertson et al., 2016). Similarly, heterologous expression of *C. intestinalis* AOX enables *in vivo* KCN tolerance of *Drosophila* (*Sophophora*) *melanogaster* Meigen, 1830 (e.g. Fernandez-Ayala at al., 2009) and mice, observed also for the isolated mitochondria (e.g. El-Khoury et al., 2013; Szibor et al., 2017) However, in mouse and *D*. (*S*.) *melanogaster* mitochondria the enzyme has been shown to become enzymatically active only when the respiratory chain cytochrome pathway is inhibited beyond coenzyme Q (Kemppainen, 2014; Rajendran et al., 2019; Szibor et al., 2017, 2020) whereas in the case of *S. cerevisiae* mitochondria the activity has been observed without the cytochrome pathway inhibition (Robertson et al., 2016). The data available for *C. gigas* AOX monitored in the oyster isolated mitochondria does not allow for conclusion concerning the suggested mechanisms (Sussarellu et al., 2013). Nevertheless, the data on *C. gigas* AOX expressed in *S. cerevisiae* mitochondria (Robertson et al., 2016) corresponds to *M. inceptum* AOX based respiration of *S. cerevisiae* cells (Fig. 2). The difference in regulation of animal AOX in *S. cerevisiae* and animal mitochondria can be explained by differences in organization of the animal and *S. cerevisiae* respiratory chain, including the complex I which precedes coenzyme Q in the animal chain but is absent in *S. cerevisiae* mitochondria (de Vries and Marres, 1987). Accordingly, it has recently been shown that in the absence of the respiratory chain cytochrome pathway inhibitors, respiratory activity of complex II, but not complex I, both providing electrons to coenzyme Q, is a pre-requisite for the engagement of AOX activity (Szibor et al., 2020).

The engagement could result in coenzyme Q oxidation and consequently diminution of ROS generation providing damages to DNA, proteins and phospholipids (Hansen et al., 2006; Srinivasan and Avadhani, 2012; Jönsson, 2019; Saari et al., 2019). Thus, the enzyme may be considered as an important element of protection against ROS-mediated oxidative stress indicated for anhydrobiosis. The oxidative stress can occur during tun formation (dehydration) and tun rehydration (Rebecchi, 2013; Rizzo et al., 2015; Schill and Hengherr, 2018) due to different mechanisms. During dehydration the oxidative stress may result from gradual elimination of water and deprivation of appropriate oxygen availability (e.g. Rebecchi, 2013) whereas during rehydration may be triggered by reoxygenation (Hermes-Lima and Storey, 1998) and reverse electron transport (Murphy, 2009; (e.g. McDonald and Gospodaryov, 2019). Moreover, it is commonly suggested that the decline in revival with increasing time in the tun stage may be caused by the oxidative stress based on non-enzymatic and/or enzymatic reactions (e.g. Guidetti and Jönsson, 2002; Rebecchi et al., 2013; Schill and Hengherr, 2018). Accordingly, it has been observed for *M. inceptum* that the amount of the oxidative DNA damage increases with the duration of the tun stage (Neumann et al., 2009) Thus, if the tardigrade AOX indeed contributes to protection against oxidative stress, the protection should be distinctly impaired in the presence of AOX inhibitor, e.g. BHAM.

The estimation of return to full activity of the anhydrobiotic *M. inceptum* appears to support important role of AOX during dehydration as the presence of BHAM during tun formation imposed distinct effect on return to full activity (Fig. 4) whereas no statistically significant effect was observed for the presence of the AOX inhibitor during tun rehydration (Fig. 5). Moreover, tun formation in the presence of BHAM enhanced the well-known effect of the tun stage duration on anhydrobiotic tardigrade recovery to full activity (e.g. Wright, 1989; Rebecchi et al., 2007; Schill and Hengherr, 2018). We observed that duration of the tun stage as well as the presence of BHAM during tun formation imposed additive effect on anhydrobiotic animal return to full activity in BHAM concentration dependent way. Therefore, it may be assumed that AOX contributes to antioxidative protection systems underlying dehydration tolerance. Thus, AOX inhibition could impair the protection and consequently the tardigrade revival.

However, the question arises whether the AOX contribution is more pronounced during dehydration or during the tun stage itself. Taking into account that the presence of BHAM, independently of the applied concentration, does not impair the average number of formed tunes (Fig. 3), one could assume that the activity of AOX is important during the tun stage. Consequently, AOX impairment caused by its inhibitor would result in additional impairment of revival after the tun stage illustrated by the delay in return to full activity and/or decreased final return to full activity. Since we observed such an effect of the BHAM presence being the inhibitor concentration-dependent and the tun stage duration-dependent we could assume that AOX contributes to *M. inceptum* tun revival and the AOX contribution increases with the tun stage duration. As mentioned above, the duration of the tun stage correlates with the possibility of oxidative damage occurrence (Neumann et al., 2009).

The appearing role of AOX in longer-term anhydrobiosis can be explained by its commonly mentioned anti-oxidative effect. Nevertheless, it should be remembered that BHAM itself may also display such an activity. However, our data indicates that the effect observed for BHAM differ distinctly from that imposed by MitoTEMPO, a well-known mitochondria-specific superoxide scavenger (Fig. 6-7). This, in turn, implies that AOX contribution is much more complex. On the one hand, AOX may ensure efficient ATP synthesis under constrained activity of the respiratory chain cytochrome pathway. Interestingly, the possibility of ATP synthesis appears to be pertinent for successful aestivation (Kayes et al., 2009), an important animal survival strategy which like anhydrobiosis requires reversible transitions to and from hypometabolic states (Storey and Storey, 2004). On the other hand, it is proposed that AOX contributes to the control of ROS generation by the mitochondrial respiratory chain modulation, also in invertebrate cells (e.g. Fernandez-Ayala et al., 2009; Letendre et al., 2012).

## CONCLUSION

The presence of functional *AOX* in *M. inceptum*, formely known as *M. tardigradum* and regarded as one of the most stress resistant tardigrade species, makes this protein a strong candidate in searching of mitochondrial proteins important for successful anhydrobiosis. Accordingly, the obtained data indicates that AOX activity is crucial for successful revival of the tun stage, particularly for longer-term anhydrobiosis and the involved mechanism is more complex than simple ROS elimination. Thus, the obtained results confirms the importance of AOX research in explanation of the role of mitochondria in the successful anhydrobiosis that may provide applicative after-effects concerning medicine, biotechnology and even space travels.

## Supporting information

**S1 Fig. Integration of the AOX gene into the yeast genome**. Plasmid pML104 expresses Cas9 endonuclease and designed sgRNA, which allows DNA cleavage in the *GAL1* region (a). The repair DNA containing the Mi-AOX gene and the GAL1 gene flanks (intergenic regions - *iGAL1*) are incorporated into the yeast genome by homologous recombination (b). Replacement of DNA fragments removes the Cas9 cleavage site and allows expression of the Mi-AOX gene under the control of the *GAL1* promoter (c). Mi, *Milnesium inceptum*.(pdf)

**S1 Data. Contains additional data concerning Fig 1A–1B**. (pdf)

## Acknowledgements

The studies were performed partly in the framework of activities of BARg (Biodiversity and Astrobiology Research group). We would like to thank Edyta Fiałkowska (Laboratory of Aquatic Ecosystems Group, Institute of Environmental Sciences, Jagiellonian University in Kraków, Poland) for the rotifer *Lecane inermis* cultures, Pushpalata Kayastha (Department of Animal Taxonomy and Ecology, Institute of Environmental Sciences, Adam Mickiewicz University, Poznań, Poland) for help in identification of the applied tardigrade species and Felix Bemm (Max Planck Institute for Developmental Biology, Tübingen, Germany) for data concerning *M. inceptum* AOX encoding gene.

