## Supplementary Figure 1 for "Mitochondrial alternative oxidase contributes to successful tardigrade anhydrobiosis"

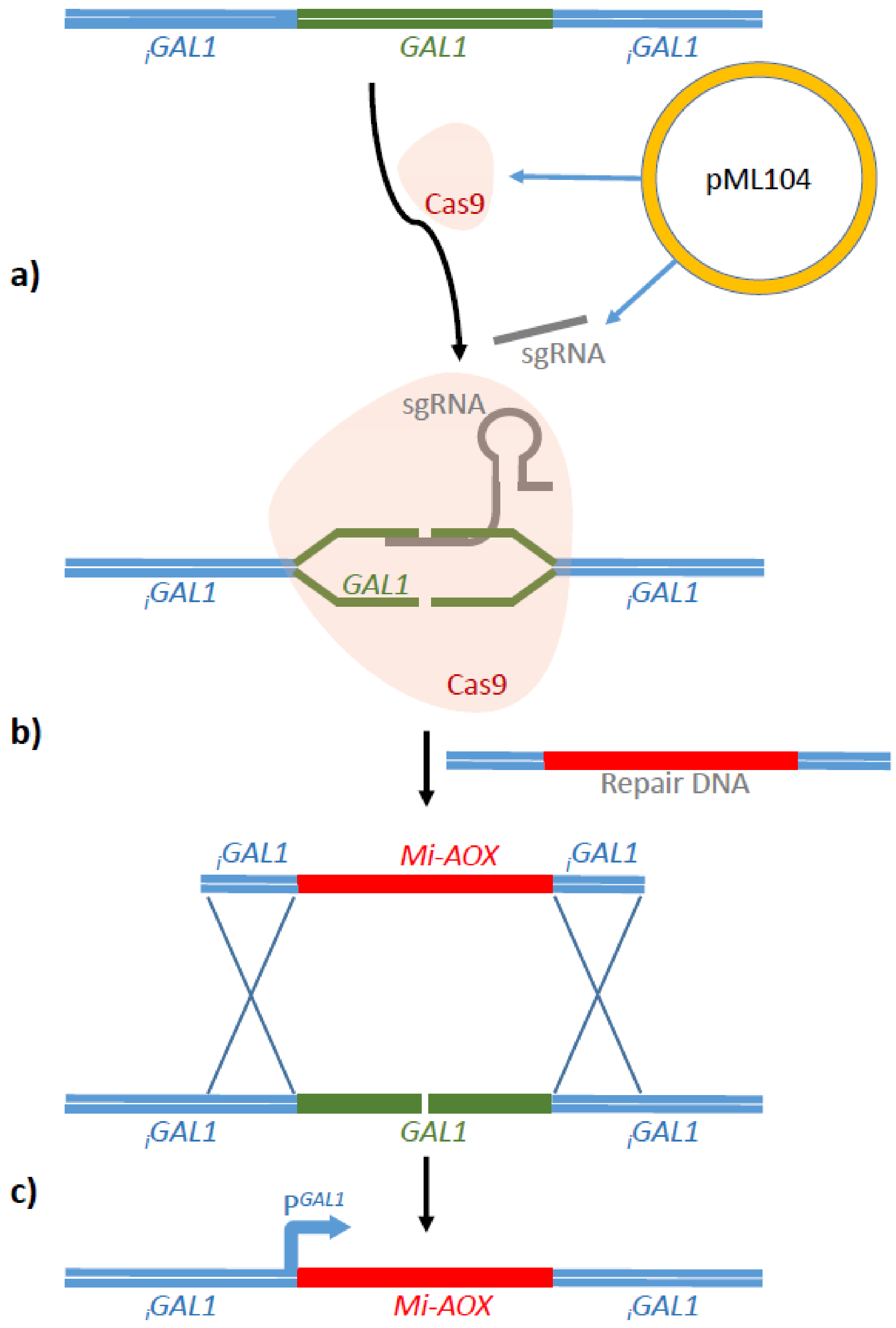


**S1 Fig.** Integration of the AOX gene into the yeast genome. Plasmid pML104 expresses Cas9 endonucleases and designed sgRNA, which allows DNA cleavage in the *GAL1* region (a). The repair DNA containing the Mt-AOX gene and the *GAL1* gene flanks (intergenic regions - *iGAL1*) are incorporated into the yeast genome by homologous recombination (b). Replacement of DNA fragments removes the Cas9 cleavage site and allows expression of the Mt-AOX gene under the control of the GAL1 promoter (c).

**Supplementary Data (S1 Data)**

Structural motives

MotifFinder analysis:

AOX, Alternative oxidase, ferritin-like diiron-binding domain.

| | |
|---|---|
| Position | 238..403 |
| Alignment<br>Query<br>Database | EAKWVRRVVFLETVAGVPGMMGAMARHLKSLRSMRKDLGWIHTLLEEAENERMHL<br>MCALELKQPTWLFKLGTVITQGIFTNVFFILYLMSPRFCHRFVGYLEEEAVRTYTKLLYDI<br>DHGSMAHWQTQPAPEIAVKYWQLPASATCRDMVLAIRADEAHHRLVNHTLEDRWL<br>ARFIFLETVARVPGMVAGMLLHLYSLRGMWRDGGWIKTLLEEAENERMHLLIFEELGG<br>PGWWFRRFVAQHQAVFYNAYFLLYLISPRLAHRFVGYLEEEAVDTYTEFLKDIEEG---<br>LKPDLPAPEIAIEYYRLGEDATLYDVFVAIRADEAEHRKVNHAC |
| Score | 260 |
| E-value | 5e-85 |

ferritin-like domain ($Fe^{2+}$ – binding domain) is marked in red

```
MPVKPLNFRGIALRRSPIASTQISYHSRPHLRSFTSNGFTHLRPLQTWQSRTPFIYDTLH
NKSTYAILGRRGLASKPQSSESSSDLDKKIADPLSDEPSKSKGKATSVSADGRQPQQQRQ
DMNEAELAKSIYGKIAAEDLTHFKREAEGTSAADDKQREKELESVVIARSVNEDKLKVGD
ELAVSLPHPVWTAESLNNVEINHLKPENKIDWLALWSVRLLRFGFDVTSGYIFGRLNEAK
WVRRVVFLETVAGVPGMMGAMARHLKSLRSMRKDLGWIHTLLEEAENERMHLMCALELKQ
PTWLFKLGTVITQGIFTNVFFILYLMSPRFCHRFVGYLEEEAVRTYTKLLYDIDHGSMAH
WQTQPAPEIAVKYWQLPASATCRDMVLAIRADEAHHRLVNHTLGSLQPGKGNPFEPGR
```

**Cellular localization**

DeepLoc-1.0 analysis:

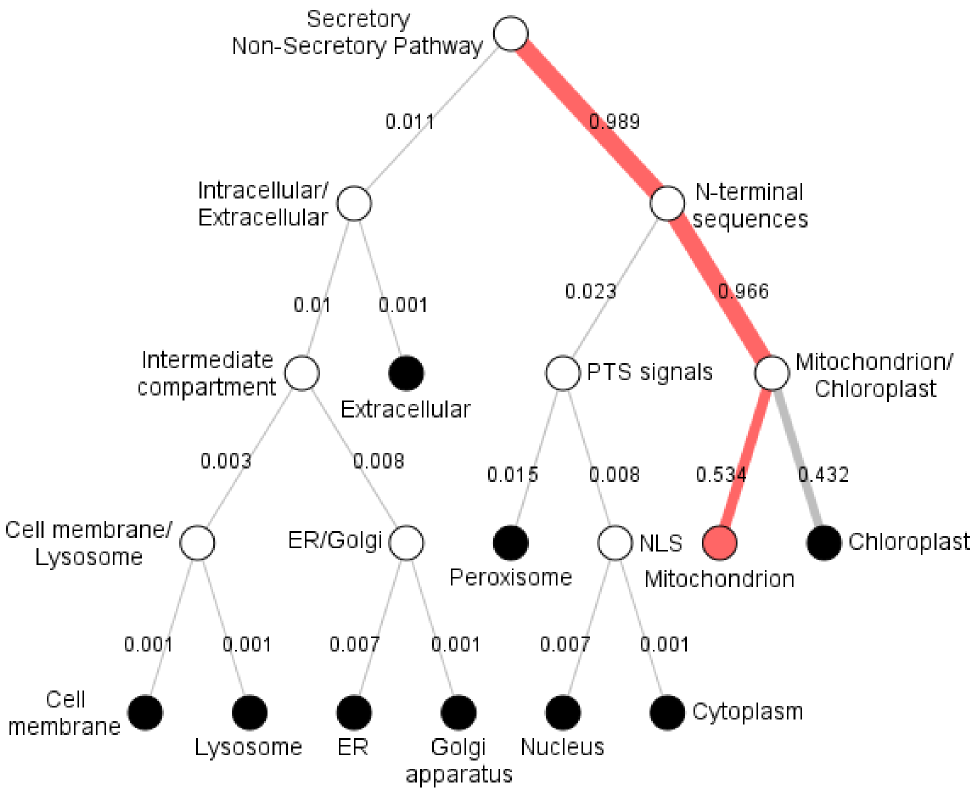


Prediction: Mitochondrion, Inner Membrane

**<u>The root-mean-square deviation of atomic positions (RMSD) indicating average distances between atoms of *M. inceptum* and *T. brucei* in their superimposed three-dimensional structures.</u>**

```
mAOX  157 Q R E K E L E S V V I A R S V N E D K L K V G D E L A V S L P H P V W T A E S L N N V E I N H L K P E N K I D W L A L W S V R L L 221(418)
tAOX    1 P V W G H T Q L N R L S F L E T V P V V P L R V S D E S S E D R P T W S L P D I E N V A I T H K K P N G L V D T L A Y R S V R T C 65(267)
RMSD      0 3 5 1 0 0 0 0 0 0 0 0 0 0 0 0 0 0 0 0 0 0 0 0 0 0 0 0 0 0 0 0 0 0 0 0 0 0 0 0 0 0 0 0 0 0 0 0 0 0 0 1 0 0 0 0 0 0 0 0 0 0 0 0 0

mAOX  222 R F G F D V T S G Y I F G R L N E A K W V R R V V F L E T V A G V P G M M G A M A R H L K S L R S M R K D L G W I H T L L E E A E 286(418)
tAOX   66 R W L F D T F S L Y R F G S I T E S K V I S R C L F L E T V A G V P G M V G G M L R H L S S L R Y M T R D K G W I N T L L V E A E 130(267)
RMSD      0 0 0 0 0 0 0 0 0 0 1 1 0 1 1 0 0 0 0 0 0 0 0 0 0 0 0 0 0 0 0 0 0 0 0 0 0 0 0 0 0 0 0 0 0 0 0 0 0 0 0 0 0 0 0 0 0 0 0 0 0 0 0 0 0

mAOX  287 N E R M H L M C A L E L K Q P T W L F K L G T V I T Q G I F T N V F F I L Y L M S P R F C H R F V G Y L E E E A V R T Y T K L L Y 351(418)
tAOX  131 N E R M H L M T F I E L R Q P G L P L R V S I I I T Q A I M Y L F L L V A Y V I S P R F V H R F V G Y L E E E A V I T Y T G V M R 195(267)
RMSD      0 0 0 0 0 0 0 0 0 0 0 0 0 0 0 0 0 0 0 0 0 0 0 0 0 0 0 0 0 0 0 0 0 0 0 0 0 0 0 0 0 0 0 0 0 0 0 0 0 0 0 0 0 0 0 0 0 0 0 0 0 0 0 0 0

mAOX  352 D I D H G S M A H W Q T Q P A P E I A V K Y W Q L P A S A T C R D M V L A I R A D E A H H R L V N H T L G S L - - - - Q P G K G N 412(418)
tAOX  196 A I D E G R L R P T - K N D V P E V A R V Y W N L S K N A T F R D L I N V I R A D E A E H R V V N H T F A D M H E K R L Q N S V N 259(267)
RMSD      0 0 0 0 0 0 0 0 0 1   1 0 1 0 0 0 0 0 0 0 0 0 0 0 0 0 0 0 0 0 0 0 0 0 0 0 0 0 0 0 0 0 0 0 0 0 0 0 0 0 0         3 0 0 0 0 0

mAOX  413 P F E P G R 418(418)
tAOX  260 P F V V L K 265(267)
RMSD      0 0 0 0 1 1
```
